# GREMI: an Explainable Multi-omics Integration Framework for Enhanced Disease Prediction and Module Identification

**DOI:** 10.1101/2023.03.19.533326

**Authors:** Hong Liang, Haoran Luo, Zhiling Sang, Miao Jia, Xiaohan Jiang, Zheng Wang, Xiaohui Yao, Shan Cong

## Abstract

Multi-omics integration has demonstrated promising performance in complex disease prediction. However, existing research typically focuses on maximizing prediction accuracy, while often neglecting the essential task of discovering meaningful biomarkers. This issue is particularly important in biomedicine, as molecules often interact rather than function individually to influence disease outcomes. To this end, we propose a two-phase framework named GREMI to assist multi-omics classification and explanation. In the prediction phase, we propose to improve prediction performance by employing a graph attention architecture on sample-wise co-functional networks to incorporate biomolecular interaction information for enhanced feature representation, followed by the integration of a joint-late mixed strategy and the true-class-probability block to adaptively evaluate classification confidence at both feature and omics levels. In the interpretation phase, we propose a multi-view approach to explain disease outcomes from the interaction module perspective, providing a more intuitive understanding and biomedical rationale. We incorporate Monte Carlo tree search (MCTS) to explore local-view subgraphs and pinpoint modules that highly contribute to disease characterization from the global-view. Extensive experiments demonstrate that the proposed framework outperforms state-of-the-art methods in seven different classification tasks, and our model effectively addresses data mutual interference when the number of omics types increases. We further illustrate the functional- and disease-relevance of the identified modules, as well as validate the classification performance of discovered modules using an independent cohort. Code and data are available at https://github.com/Yaolab-fantastic/GREMI.

## I. Introduction

**C**OMPLEX diseases often present a multifactorial nature [1]–[3], which has motivated recent advancements in high-throughput omics data acquisition, and further stimulated the growth of multi-omics integration as a rapidly expanding research field. Compared to single-omics studies [4], [5], multi-omics studies enable the capture of complementary information across various molecular layers such as DNA, RNA, and protein, therefore improving the understanding of underlying biological processes and molecular mechanisms involved in complex diseases [6], [7].

However, as the volume and variety of omics data continue to increase, multi-omics research faces significant challenges due to the mutual interference both within individual omics and between different types of omics data [8]–[10]. Moreover, the complexity of omics data poses a significant issue in terms of explanation. The high dimensionality and intricate interconnections between various data types make it difficult to discern meaningful patterns and relationships.

Artificial neural networks have been employed to tackle the problem of efficiently learning low-dimensional representations from high-dimensional omics data [11]–[13] (e.g., autoencoder [14], [15], variational autoencoder [16], generative adversarial networks (GAN) [17], [18], and graph neural networks (GNN) [19]). Among these, GNN exhibits substantial potential due to the natural advantage of graph architecture for capturing and passing interaction information between molecules. For example, Mogonet [19] employed graph convolutional networks (GCN) for disease prediction, by constructing sample similarity graphs with nodes denoting samples and edges denoting the sample similarities. *Despite the innovative utilization of deep graph architecture in multi-omics studies, such sample similarity networks cannot fully capture the comprehensive information inherent in molecules*. Alternatively, a number of studies integrated general protein-protein interaction networks (e.g., HINT [20], STRING [21], [22]) into disease prediction to account for the co-functionality of multiple genes in biological processes. Recently, transcriptomics studies highlighted the superiority of disease-specific co-expression networks for diagnosis classification. For example, Ramirez et al. [23] applied Spearman correlation to construct a co-expression network and employed GCNs for cancer classifications. Xing et al. [24] built the co-expression network and showed the performance of single-omics data on disease prediction using the attention-based GNN model.

Besides learning omics-specific representations, efforts increasingly focused on developing integration strategies to harness the correlated and/or complementary information implicated across multiple omics data. The integration strategy can generally be categorized into three types: early, joint, and late integration. Early integration involves merging various omics sources before feeding them into a model. While this method is straightforward, it can suffer from increased dimensionality and noise as well as overlook the specific distribution of each omics type [25]. Late integration aggregates prediction results produced by each omics (e.g., [19]). While it adapts well to each individual omics, late fusion cannot fully consider the cross-modal interactions [25]. Recent studies indicated that joint strategy, which combined representations from various omics, offered enhanced classification accuracy [25]–[27]. Furthermore, Han et al. [26] introduced confidence-based fusion into joint integration, to promote the prediction performance by quantifying the classification certainty of each omics.

Furthermore, explanations bear significant importance in biomedicine, yet current efforts have paid insufficient attention to this critical issue of disease prediction - **the identification of meaningful biomarkers**. Current research on biomarker discovery typically adopts a simple ablation method to evaluate the contribution of each feature in terms of classification accuracy, from either the global or local view [12], [28]. Global explanation is the most commonly used strategy that assesses the importance of features across the entire sample set (e.g., [19], [24], [26]). The local strategy focuses on offering insights into the predictive mechanisms of individual samples and is also starting to be used in omics studies. For example, the shapley additive explanations (SHAP) method was used to quantify the discriminative ability of each gene in distinguishing disease subgroups [29]. *However, whether employing the global or local strategy, existing research primarily focused on identifying individual biomarkers while overlooking the co-functional information inherent in multiple molecules*. Functional elements often interact with each other, rather than independently exert effects on disease. The intricate interplay of factors in complex diseases, combined with the advancements in graph modeling and subgraph-based interpretation [30], drives our pursuit to identify biomarkers from a co-functional module perspective.

Based on the above observations, we propose a framework, named **GR**aph-based **E**xplainable **M**ulti-omics **I**ntegration (GREMI), for disease prediction and biomarker discovery. The proposed model leverages graph attention networks and a confidence-adaptive mechanism to capture both within- and across-omics information. Then a subgraph-based interpreter is designed to promote the model explainability by extracting functional modules that contribute to classification. We demonstrate the superior performance of our model in terms of both classification and module-level biomarker identification, utilizing four benchmark and four in-house datasets. Our main contributions can be summarized as follows:

- We employ a graph attention architecture to better capture within-omics interactions and generate sample-wise latent features, and utilize a joint-late mixed integration strategy with true class probability to adaptively balance feature-level and omics-level contributions, thereby improving learning and predictive performance.
- We introduce module-level interpretative mechanisms, adopting Monte Carlo tree search to probe subgraphs and pinpoint disease modules from a local-global combined view. This subgraph-based strategy is more intuitive, considering that complex diseases often result from the interplay of multiple dysfunctional molecules.
- Our proposed model significantly surpasses the state-of-the-art methods across seven classification tasks, underscoring the generalizability of GREMI.
- We validate the identified biomarker modules in an independent cohort, emphasizing the capability of our framework for disease target discovery.

## II. Materials and methods

The overview of the proposed method is illustrated in Fig. 1, which comprises two main phases: the prediction phase (Fig. 1(a)) and the interpretation phase (Fig. 1(b)). During the prediction phase, co-functional networks tailored to each omics were constructed for each sample, followed by multi-level graph attention networks (GAT) to produce graph representations. Subsequently, the true class probability was implemented to estimate feature-level confidences, with joint fusion adopted to incorporate inter-omics information for final prediction. In the interpretation phase, the Monte Carlo tree search (MCTS) algorithm was used to locally explore subgraphs (i.e., sample-wise candidate modules), followed by a rank-based evaluation mechanism to extract globally important ones (i.e., consensus modules). Permutation and functional annotation were executed to evaluate the performance of identified modules.

**Fig. 1.**
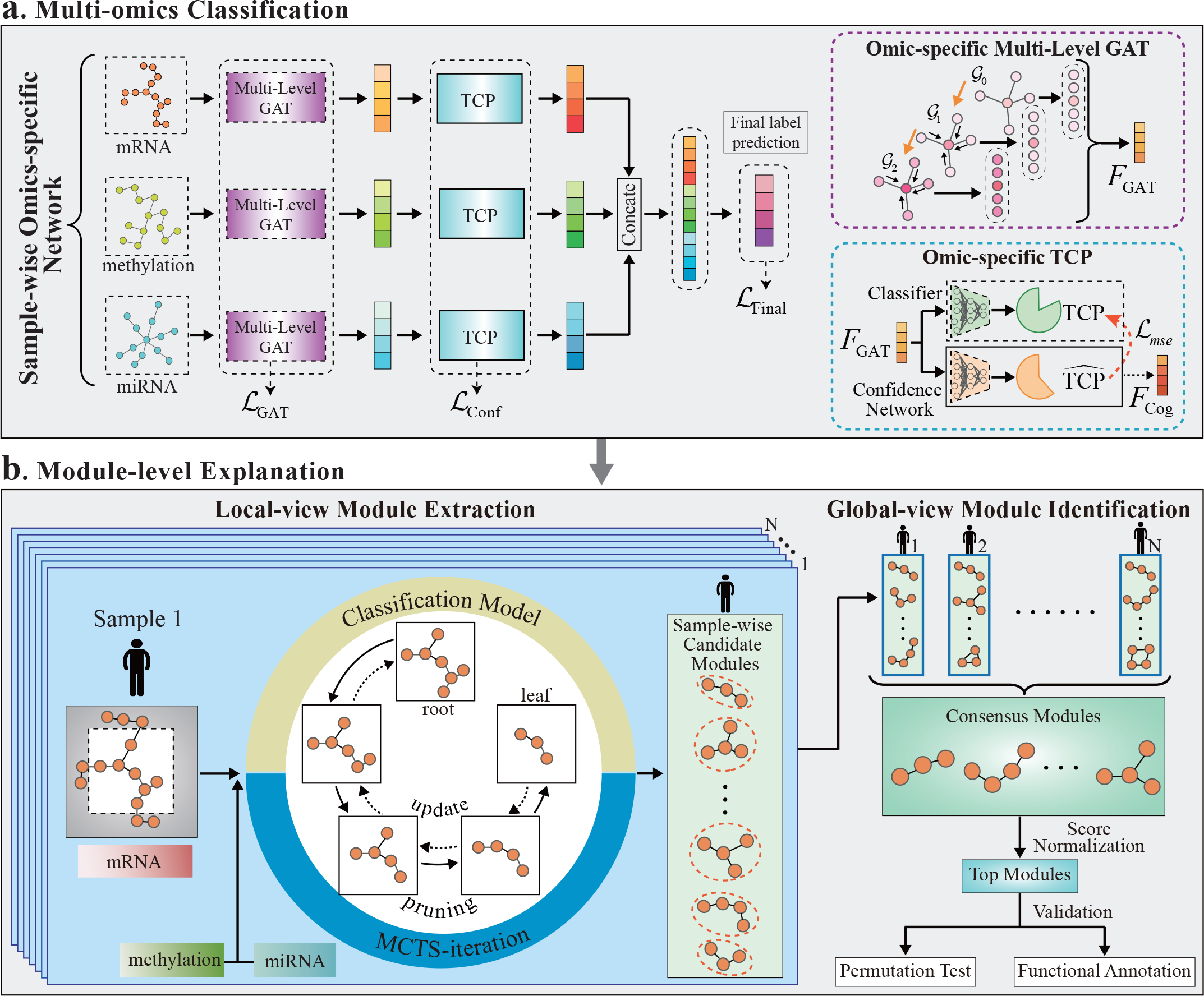
Framework overview. (a) **Prediction phase:** For each sample, omics-specific co-functional networks are constructed using the WGCNA; Multi-level GAT is used to capture functional interactions within each omics and produce omics-specific graph representations; TCP is implemented to evaluate feature-level classification confidences and optimize omics-specific representations; Joint fusion concatenates confidence-adapted features to incorporate inter-omics information and makes final predictions. (b) **Interpretation phase:** With the trained model and co-functional networks as inputs, MCTS is used to generate important subgraphs as local interpretations of each sample; Consensus modules across samples are extracted and ranked based on their normalized importance scores; Permutation and functional annotation are performed to evaluate the reported modules.

### A. Multi-omics datasets and co-functional network construction

#### 1) Multi-omics datasets

To demonstrate the efficacy of our proposed model, we conducted extensive experiments on eight public datasets (Table I): four benchmark datasets including ROSMAP for Alzheimer’s disease (AD) classification, BRCA for PAM50-defined breast cancer subtype classification, LGG for low-grade glioma (LGG) grade 2 vs. grade 3 classification, and KIPAN for renal cell carcinoma subtype classification; three in-house processed datasets including LUAD for lung adenocarcinoma subtype classification, THCA for thyroid cancer subtype classification, and UCEC for uterine corpus endometrial carcinoma grade classification; and one validation dataset LGG-V (LGG grade 2 vs. grade 3) obtained from an independent cohort, the Chinese Glioma Genome Atlas (CGGA, http://www.cgga.org.cn/), to evaluate the performance of modules identified from the LGG. We also investigated how varying the number of features influenced model predictions on LGG and KIPAN by using the top 1,000 (if available) and top 200 features based on the ANOVA *F* -value. Detailed information regarding data acquisition and preprocessing is available in Supplementary Methods.

**TABLE I.** Summary of datasets used in this study.

| Dataset | Type | Source | Sample | Number of raw features<sup>a</sup> |  |  | Number of features for training<sup>b</sup> |  |  |
| --- | --- | --- | --- | --- | --- | --- | --- | --- | --- |
|  |  |  |  | mRNA | methy | miRNA | mRNA | methy | miRNA |
| ROSMAP | Benchmark | AMPAD | NC: 169, AD: 182 | 55,889 | 23,788 | 309 | 200 | 200 | 200 |
| BRCA | Benchmark | TCGA | Luminal A: 436, Luminal B: 147, HER2-enriched: 46, Normal-like: 115, Basal-like: 131 | 20,531 | 20,106 | 503 | 1,000 | 1,000 | 503 |
| LGG | Benchmark | TCGA | Grade 2: 246, Grade 3: 264 | 20,531 | 20,114 | 548 | 2,000<br>1,000<br>200 | 2,000<br>1,000<br>200 | 548<br>548<br>200 |
| KIPAN | Benchmark | TCGA | KICH: 66, KIRC: 318, KIRP: 274 | 20,531 | 20,111 | 445 | 2,000<br>1,000<br>200 | 2,000<br>1,000<br>200 | 445<br>445<br>200 |
| LUAD | In-house | TCGA | Mixed: 90, Not otherwise specified: 281 | 20,531 | 20,115 | 529 | 200 | 200 | 200 |
| THCA | In-house | TCGA | Follicular: 102, Classical/usual: 356 | 20,531 | 20,108 | 517 | 200 | 200 | 200 |
| UCEC | In-house | TCGA | Grade 2: 122, Grade 3: 316 | 20,531 | 20,108 | 554 | 200 | 200 | 200 |
| LGG-V | Validation | CGGA | Grade 2: 188, Grade 3: 255 | 23,987 | 14,476 | 829 | 5 | 0 | 0 |

<sup>a</sup> lists the number of features obtained from the original datasets. <sup>b</sup> lists the number of features used for model training.

#### 2) Sample-wise co-functional network construction

Functional interaction network holds significance for elucidating the pathogenesis of complex diseases. To harness the inherent co-functional information between omics molecules, we initially employed weighted correlation network analysis (WGCNA [31]) to establish omics-specific co-functional networks (Fig. 1(a)). Considering mRNA as an example, the process of obtaining sample-wise graphs comprised the following steps: 1) an adjacency matrix was constructed from WGCNA, where each element represented the strength of expression correlation between each pair of mRNAs; 2) an edge matrix was obtained by thresholding the adjacency matrix; 3) for each individual sample, a co-expression network was devised by assigning each mRNA expression values to corresponding nodes as node features. Similarly, we formulated the sample-wise co-methylation and miRNA co-expression graphs for each dataset.

### B. GATs for omics-specific representation learning

GNNs facilitate the utilization of interactive relationships contained within omics data, among which, the graph attention mechanism is more capable of adaptively aggregating neighbor information [32]. Xing et al. [24] demonstrated the superiority of GAT in a single-omics study. To this end, we employed GATs for omics-specific graph representation learning, where sample-wise GATs were trained for each omics type, as illustrated in Fig. 1(a). Specifically, multi-head multi-level GATs were implemented to represent sample-wise co-functional networks. Illustrated with mRNA data, each sample-wise co-expression network could be viewed as a graph *G*_0_ = (**X, E**), where **X** *∈*R^*n×d*^ is the feature matrix, **E** *∈*R^*n×n*^ is the edge matrix, *n* is the number of nodes and *d* is the number of features (here *d* = 1). Taking *G*_0_ as input, a GAT could be built by stacking graph attention layers with multi-head attention. Each layer is defined as:

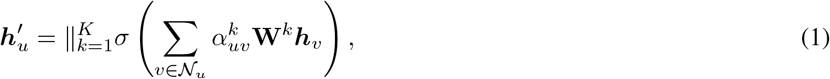

where ∥ represents concatenation of *K* heads, ***h***_*v*_ is the input features of node *v*, 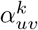 is the *k*th normalized attention coefficients, **W**^*k*^ is the weight matrix of head *k, N*_*u*_ is the first-order neighbors of node *u*, and *σ*(·) is a nonlinear activation function. *α*_*uv*_ is calculated from the attention mechanism *a*:

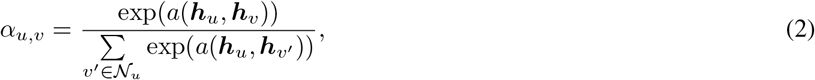

with *a*(***h***_*u*_, ***h***_*v*_) representing the importance of node *v* in relation to node *u*.

In addition to aggregating the node-level molecular features, we further adopted a multi-level approach to promote the information aggregation of molecular modules. In particular, a higher-level graph *G*_1_ was generated by applying multi-head GAT layers on *G*_0_, and *G*_2_ was derived from *G*_1_ in a similar manner. After that, graph embeddings from three levels were concatenated, generating more enriched multi-level representations. These representations were subsequently input into fully-connected layers to produce omics-specific embeddings, denoted as *F*_GAT_. Meanwhile, for each omics type, the GAT classifier was trained to incorporate within-omics information into prediction:

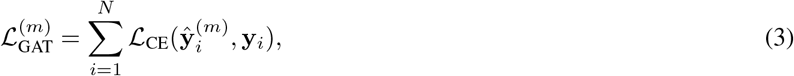

where *L*_CE_ is the cross-entropy loss function, *N* is the number of training samples, **y**_*i*_ is the true label, and 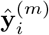 is the prediction from the *m*th omics data.

The inherent heterogeneity among multi-omics data, coupled with external influences such as differences in data acquisition and storage conditions [33], [34], presents significant challenges for data integration [10], [35]. Han et al. [26] showed the performance of true class probability in multi-modal learning. Therefore, in addition to improving the predictive capacity of each omics data, we also adopted the trustworthy strategy to evaluate and adaptively regulate the prediction confidence associated with each type of omics data (see Fig. 1(a)).

The traditional approach for confidence inference is the maximum class probability (MCP). For omics *m*, given the input feature matrix **X**^(*m*)^, the classifier can be interpreted as a probabilistic model. Utilizing the softmax function, it assigns predictive distribution *P* (*Y* |**X**^(*m*)^) for each class *k* = {1, …, *K*}. Subsequently, the predicted class can be inferred as *ŷ* = arg max_*k∈{*1,…,*K*}_ *P* (*Y* = *k* |**x**^(*m*)^). It can be observed that MCP selects the highest softmax probability, resulting in over-confidence for incorrect predictions.

To solve this problem, the true class probability (TCP) confidence criterion [36] has been proposed, aiming for separate assignments of low and high confidences for erroneous and successful predictions, respectively. In our model, we adopt the TCP criterion to obtain more reliable prediction confidences for various omics:

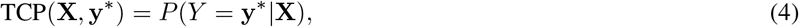

where **y**^*∗*^ is the true label vector. Equation (4) shows that the TCP and MCP are equivalent to each other when the sample is correctly classified, while the former produces a lower value for the misclassified sample.

However, the TCP confidence cannot be directly estimated on the testing set due to the unavailability of true labels. Such that, for *m*th omics dataset, a confidence neural network with parameters *θ*^(*m*)^ is introduced to estimate the TCP confidence 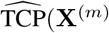, *θ*^(*m*)^) on the training data, as shown in Fig. 1(a). Specifically, the *𝓁*_2_ loss is used to train the confidence network:

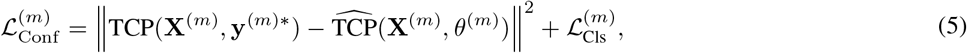

where 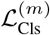 is cross-entropy loss for omics-specific classifier. In summary, the TCP module constructs an omics-specific classifier and a confidence neural network on top of the rich representations from the GAT layers and produces confidence scores for each omics data type.

### D. Multi-omics integration

Enhanced interactivity and informativeness generated by the multi-level GAT increase heterogeneity, which complicates inter-omics analyses. To address these intricacies, we adopted a joint-late-mixed integration technique, leveraging an omics-level confidence mechanism to modulate the contributions stemming from inter-omics fusion across different omics datasets. GAT encoded omics-specific representations 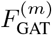 were transformed into omics informativeness, which was denoted as cognizant-level features 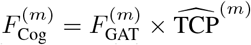. Then, we introduced a selective attention mechanism to generate more discriminative features: 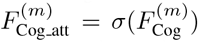, where *σ* denotes an attention activate function. This mechanism not only emphasizes salient features but also reduces the effects of non-informative attributes, focusing exclusively on active informed content. Subsequently, the features from multiple omics were concatenated for final classification. Overall, the total loss can be expressed as

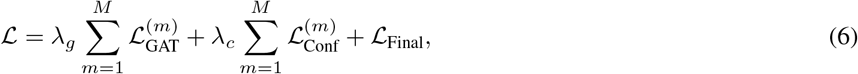

where *λ*_*g*_ and *λ*_*c*_ denote hyperparameters for adjusting different losses, and _Final_ is the cross-entropy loss for the final classification. We set both hyperparameters *λ*_*g*_ and *λ*_*c*_ equal to one across our experiments.

### E. Module-level biomarker identification and evaluation

We employed a multi-view multi-level strategy to identify biomarkers that co-functionally influenced the disease, as shown in Fig. 1(b). Taking the mRNA co-expression network as an example, we first used classification accuracy to measure the importance of each feature and obtain the feature-level importance. Features with above-average accuracy decreasing were collected to form a concentrated graph for subsequent analyses. Then we applied MCTS on to generate sample-wise important modules (i.e., local-view candidate modules). Subsequently, we extracted subgraphs that conserve high-importance scores across all samples (i.e., global-view important modules) and ranked these modules by their scores which were normalized to the number of nodes. Finally, we evaluated the top modules via permutation test and functional annotation. We described the details of MCTS-based sample-wise module identification and evaluation below.

#### 1) Local-view module extraction with MCTS

Compared to feature-level biomarkers, module-level biomarkers are of greater significance in elucidating the biological processes that underlie complex diseases. However, searching and evaluating all possible connected subgraphs from an input graph is challenging due to the huge search space, especially for graphs with hundreds or even thousands of nodes. The search problem can be efficiently solved by MCTS combined with a model-based scoring function [37].

Formally, the root of the search tree that corresponds to the input graph *G* is denoted as *S*_0_, and each child node *S*_*i*_ represents a subgraph *G*_*i*_ that is derived from a sequence of node-pruning on *G*, and the edge linking parent and child nodes represents pruning action *a*. Specifically, given a parent node *S*_*i*_ with an action *a*_*ij*_, subgraph _*j*_ is obtained by applying pruning action *a*_*ij*_ on *S*_*i*_. Then the edge (*S*_*i*_, *a*_*ij*_) records the following statistics: a visit count *N* (*S*_*i*_, *a*_*ij*_) that denotes the number of visits of action *a*_*ij*_, a total action value *W* (*S*_*i*_, *a*_*ij*_) that represents the total reward of all selected *a*_*ij*_ visits, a normalized action value *Q*(*S*_*i*_, *a*_*ij*_) = *W* (*S*_*i*_, *a*_*ij*_)*/N* (*S*_*i*_, *a*_*ij*_), and a prediction score *P* (*S*_*i*_, *a*_*ij*_) = Score(GAT(*·*),*G, G* _*j*_) that indicates the importance of subgraph *G*_*j*_ for classification.

As shown in Fig. 2, given the input graph and trained classifier, MCTS searches for subgraphs by selecting a path from the root *S*_0_ to a leaf node *S*_*l*_ iteratively over three steps:

**Fig. 2.**
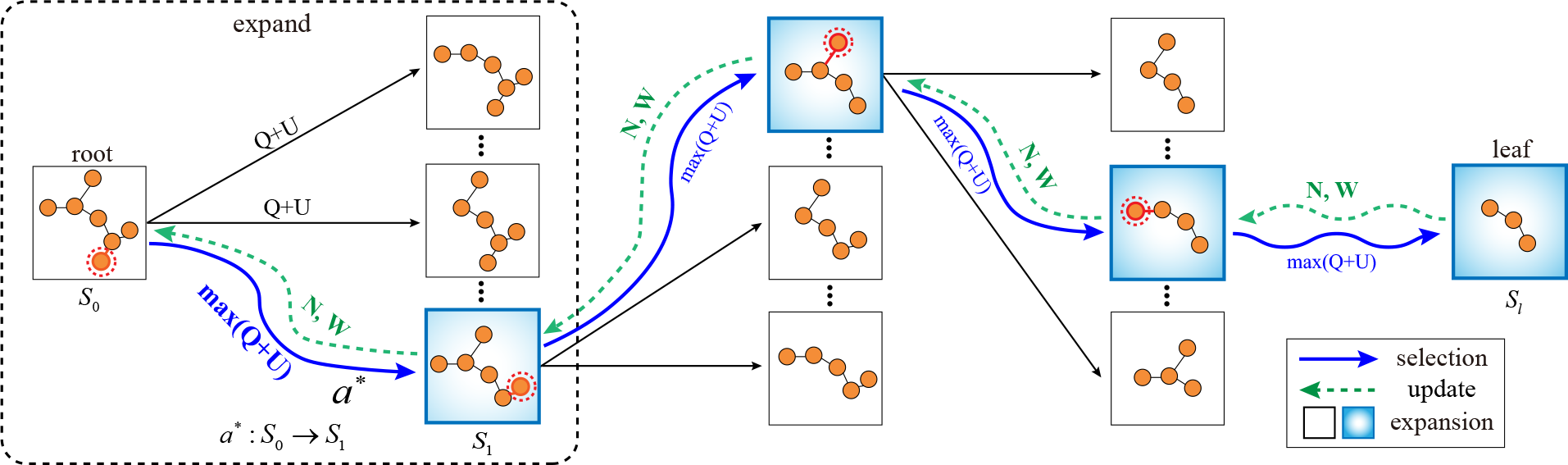
MCTS in module identification. In each iteration, the MCTS performs expansion and selection to derive a path from root node ***S***_**0**_ to leaf node ***S***_***l***_. In the expansion-selection step, subgraphs are generated by pruning the parent node, and the child node with a maximum ***Q* + *U*** value is selected (e.g., ***a***^***∗***^ **: *S***_**0**_→ ***S***_**1**_), until reaching a leaf node. Then the leaf node ***S***_***l***_ is evaluated for its prediction performance based on the trained GREMI model. After that, the count values ***N*** and action values ***W*** on the selected path are updated.

- Action selection: At each parent node *S*_*i*_, the pruning action is selected according to the following criterion:

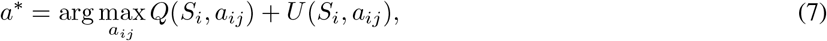

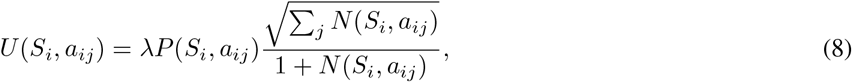

where *λ* is a hyperparameter to trade-off the exploration and exploitation.
- Subgraph evaluation: The score function is used to evaluate the prediction performance of the selected leaf node, denoted as Score(GAT(*·*), *G, G*_*l*_).
- Backward pass: Edge statistics associated with the selected path are updated:

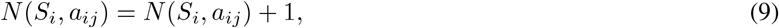

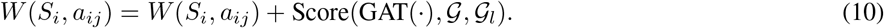

In the end, for each sample, the subgraphs from the searching paths were ranked by their prediction scores and the top findings form a candidate set of sample-wise modules. Note that we set upper and lower bounds on the size of the module to obtain results that could serve an explanatory purpose while being sufficiently concise. Similar procedures were applied to the other two omics networks and generated methylation- and miRNA-specific candidate modules for each sample.

#### 2) Global-view important module identification

Local-view candidate modules were tailored to each sample. In order to find modules that are active in the population, we further extracted the important modules from a global perspective. That is, the consensus modules which appeared consistently across the cohort as well as exhibited the highest normalized impact score were lifted out. These modules were recognized as important module-level biomarkers for the disease explanation (Fig. 1(b)).

#### 3) Permutation evaluation and annotation for identified modules

We conducted 1, 000 permutation tests to evaluate the statistical significance of identified modules. For each module, we randomly extracted 1, 000 connected subgraphs of the same size from the original network and compared their classification performance using the ablation approach. Additionally, we performed the pathway enrichment analysis using Gene Ontology (GO) and KEGG databases, to assess module functionality.

### F. Independent validation of identified modules

We utilized an independent dataset LGG-V from the CGGA [38] to validate the efficacy of the identified modules. In particular, an SVM classifier with five-fold cross-validation was employed to assess the classification performance of biomarkers reported from the LGG dataset. We additionally conducted a permutation test (N = 1, 000) to evaluate the discriminative performance of our identified biomarkers compared to the randomly selected molecules.

## III. RESULTS

We assessed the performance of our proposed model in comparison to state-of-the-art methods across various classification tasks. For the binary classification, we used accuracy (ACC), F1 score (F1), and area under the receiver operating characteristic curve (AUC). For the multiclass classification, we used ACC, average F1 score weighted (F1-weighted), and macro-averaged F1 score (F1-macro). We run experiments five times to obtain the mean and standard deviation. We further employed a t-test to show whether our model demonstrates statistically significant improvements over the state-of-the-art methods.

### A. Classification performance comparison

We evaluated the performance of our model against fourteen computational methods across four benchmark datasets, including six single-omics classifiers that employ early integration (KNN [39], SVM [40], Lasso [41], random forest (RF) [42], XGboost [43], and fully connected neural networks (NN) [44]), and seven multi-omics classifiers including group-regularized ridge regression (GRidge) [45], BPLSDA [46], block PLSDA with sparse constraints (BSPLSDA) [46], concatenation of final representations (CF) [47], gated multimodal fusion (GMU) [48], and two state-of-the-art methods (Mogonet [19] and Dynamics [26]). Moreover, we performed comparisons with Mogonet and Dynamics using three in-house processed datasets, to further demonstrate the performance of our model.

Table II and III show the comparison results on the four benchmark datasets (i.e., ROSMAP, BRCA, LGG, and KIPAN), where our model exhibits superior performance in both binary and multiclass classification tasks. Specifically, compared with the suboptimal method (i.e., Dynamics), our model achieved significant improvements in most evaluation metrics, except for the AUC of AD classification. This highlighted the informativeness of disease-specific networks and the efficacy of graph attention in information consolidation. On the other hand, our model significantly outperformed Mogonet, underscoring the advantages of employing confidence-based adaptive fusion to identify and filter informative modalities. Moreover, from Table III, we can observe that the performance of our model increases gradually as the number of features decreases. This highlights the ability of the graph neural network to represent samples using pivotal features.

**TABLE II.**
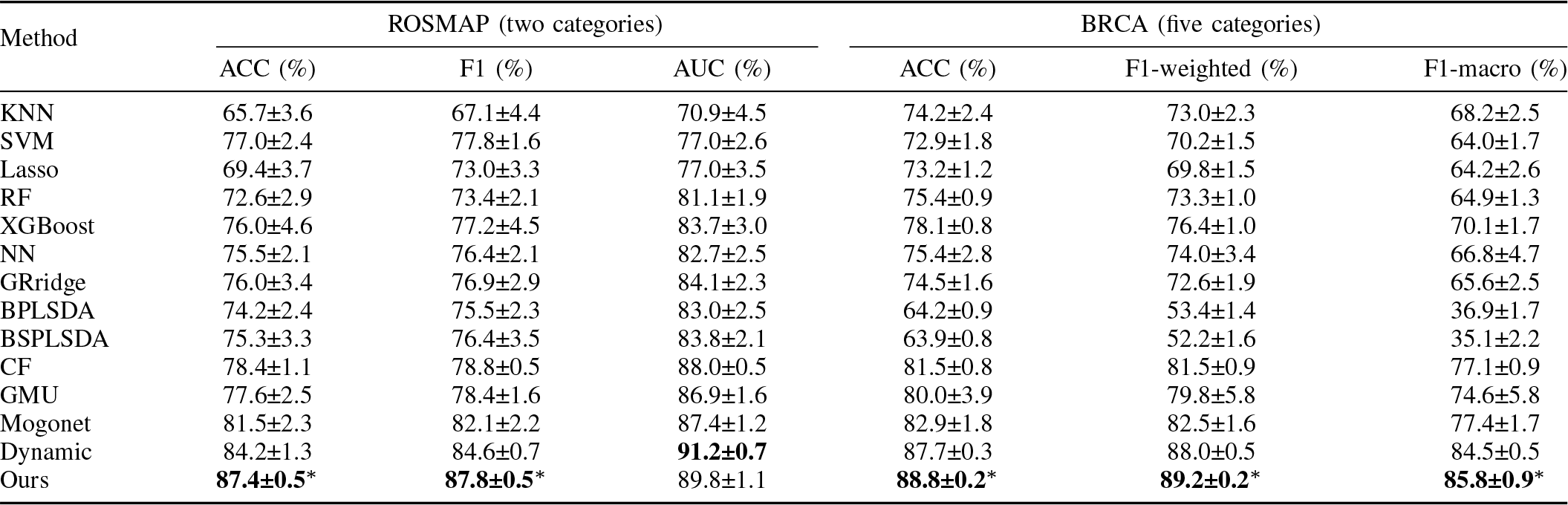
Quantitative comparison with various models on the ROSMAP and BRCA datasets. Mean and standard deviation are reported. The best results are in bold. Results with ‘*’ indicate that our model is significantly better (***P <*** 0.05) than the suboptimal method when using the independent t-test.

| Method | ROSMAP (two categories) |  |  | BRCA (five categories) |  |  |
| --- | --- | --- | --- | --- | --- | --- |
|  | ACC (%) | F1 (%) | AUC (%) | ACC (%) | F1-weighted (%) | F1-macro (%) |
| KNN | 65.7±3.6 | 67.1±4.4 | 70.9±4.5 | 74.2±2.4 | 73.0±2.3 | 68.2±2.5 |
| SVM | 77.0±2.4 | 77.8±1.6 | 77.0±2.6 | 72.9±1.8 | 70.2±1.5 | 64.0±1.7 |
| Lasso | 69.4±3.7 | 73.0±3.3 | 77.0±3.5 | 73.2±1.2 | 69.8±1.5 | 64.2±2.6 |
| RF | 72.6±2.9 | 73.4±2.1 | 81.1±1.9 | 75.4±0.9 | 73.3±1.0 | 64.9±1.3 |
| XGBoost | 76.0±4.6 | 77.2±4.5 | 83.7±3.0 | 78.1±0.8 | 76.4±1.0 | 70.1±1.7 |
| NN | 75.5±2.1 | 76.4±2.1 | 82.7±2.5 | 75.4±2.8 | 74.0±3.4 | 66.8±4.7 |
| GRidge | 76.0±3.4 | 76.9±2.9 | 84.1±2.3 | 74.5±1.6 | 72.6±1.9 | 65.6±2.5 |
| BPLSDA | 74.2±2.4 | 75.5±2.3 | 83.0±2.5 | 64.2±0.9 | 53.4±1.4 | 36.9±1.7 |
| BSPLSDA | 75.3±3.3 | 76.4±3.5 | 83.8±2.1 | 63.9±0.8 | 52.2±1.6 | 35.1±2.2 |
| CF | 78.4±1.1 | 78.8±0.5 | 88.0±0.5 | 81.5±0.8 | 81.5±0.9 | 77.1±0.9 |
| GMU | 77.6±2.5 | 78.4±1.6 | 86.9±1.6 | 80.0±3.9 | 79.8±5.8 | 74.6±5.8 |
| Mogonet | 81.5±2.3 | 82.1±2.2 | 87.4±1.2 | 82.9±1.8 | 82.5±1.6 | 77.4±1.7 |
| Dynamic | 84.2±1.3 | 84.6±0.7 | **91.2±0.7** | 87.7±0.3 | 88.0±0.5 | 84.5±0.5 |
| Ours | **87.4±0.5\*** | **87.8±0.5\*** | 89.8±1.1 | **88.8±0.2\*** | **89.2±0.2\*** | **85.8±0.9\*** |

**TABLE III.**
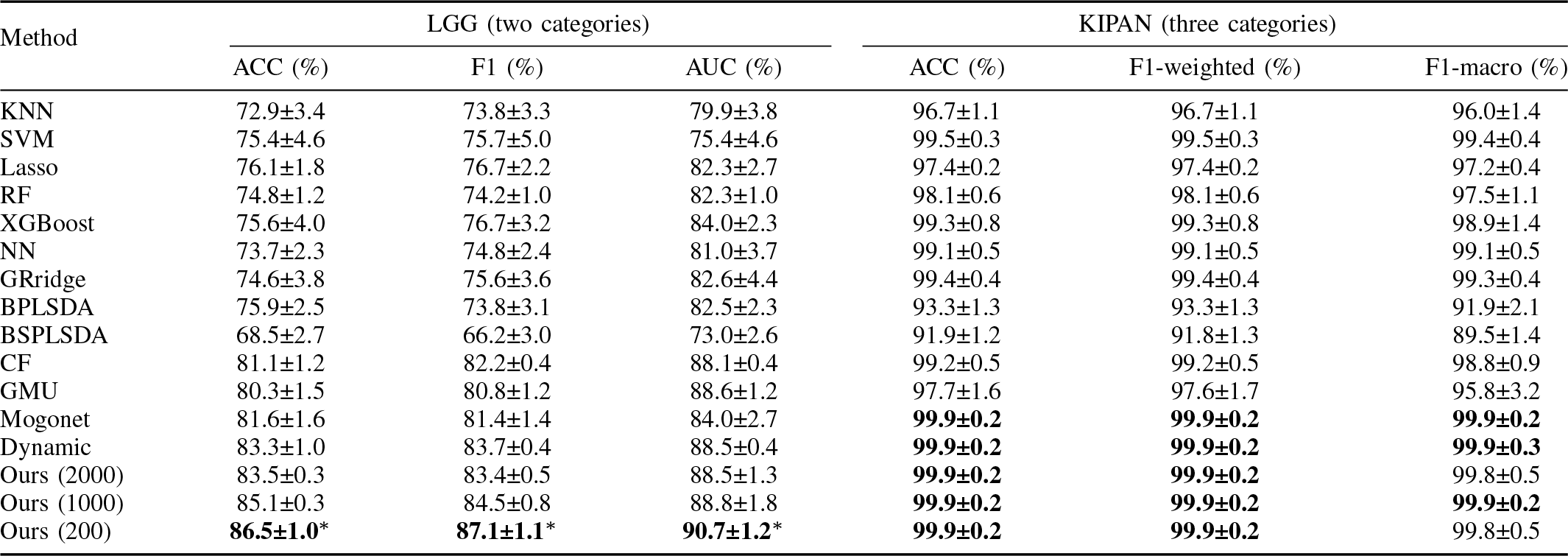
Quantitative comparison with various models on the LGG and KIPAN datasets. Mean and standard deviation are reported. The best results are in bold. Results with ‘*’ indicate that our model is significantly better (***P <*** 0.05) than the suboptimal method when using the independent t-test.

| Method | LGG (two categories) |  |  | KIPAN (three categories) |  |  |
| --- | --- | --- | --- | --- | --- | --- |
|  | ACC (%) | F1 (%) | AUC (%) | ACC (%) | F1-weighted (%) | F1-macro (%) |
| KNN | 72.9±3.4 | 73.8±3.3 | 79.9±3.8 | 96.7±1.1 | 96.7±1.1 | 96.0±1.4 |
| SVM | 75.4±4.6 | 75.7±5.0 | 75.4±4.6 | 99.5±0.3 | 99.5±0.3 | 99.4±0.4 |
| Lasso | 76.1±1.8 | 76.7±2.2 | 82.3±2.7 | 97.4±0.2 | 97.4±0.2 | 97.2±0.4 |
| RF | 74.8±1.2 | 74.2±1.0 | 82.3±1.0 | 98.1±0.6 | 98.1±0.6 | 97.5±1.1 |
| XGBoost | 75.6±4.0 | 76.7±3.2 | 84.0±2.3 | 99.3±0.8 | 99.3±0.8 | 98.9±1.4 |
| NN | 73.7±2.3 | 74.8±2.4 | 81.0±3.7 | 99.1±0.5 | 99.1±0.5 | 99.1±0.5 |
| GRidge | 74.6±3.8 | 75.6±3.6 | 82.6±4.4 | 99.4±0.4 | 99.4±0.4 | 99.3±0.4 |
| BPLSDA | 75.9±2.5 | 73.8±3.1 | 82.5±2.3 | 93.3±1.3 | 93.3±1.3 | 91.9±2.1 |
| BSPLSDA | 68.5±2.7 | 66.2±3.0 | 73.0±2.6 | 91.9±1.2 | 91.8±1.3 | 89.5±1.4 |
| CF | 81.1±1.2 | 82.2±0.4 | 88.1±0.4 | 99.2±0.5 | 99.2±0.5 | 98.8±0.9 |
| GMU | 80.3±1.5 | 80.8±1.2 | 88.6±1.2 | 97.7±1.6 | 97.6±1.7 | 95.8±3.2 |
| Mogonet | 81.6±1.6 | 81.4±1.4 | 84.0±2.7 | **99.9±0.2** | **99.9±0.2** | **99.9±0.2** |
| Dynamic | 83.3±1.0 | 83.7±0.4 | 88.5±0.4 | **99.9±0.2** | **99.9±0.2** | **99.9±0.3** |
| Ours (2000) | 83.5±0.3 | 83.4±0.5 | 88.5±1.3 | **99.9±0.2** | **99.9±0.2** | 99.8±0.5 |
| Ours (1000) | 85.1±0.3 | 84.5±0.8 | 88.8±1.8 | **99.9±0.2** | **99.9±0.2** | **99.9±0.2** |
| Ours (200) | **86.5±1.0\*** | **87.1±1.1\*** | **90.7±1.2\*** | **99.9±0.2** | **99.9±0.2** | 99.8±0.5 |

We further compared our model with two state-of-the-art methods using three in-house datasets. Table IV presents that our model yields the best performance across three datasets. The Dynamics method underperformed on certain datasets, possibly due to the inability of the MLP encoder to capture intra-omics interaction information sufficiently. Although Mogonet outperformed Dynamics, our model surpassed Mogonet, highlighting that sample-wise molecular interaction networks contain more valuable information than the sample-sample similarity network.

**TABLE IV.**
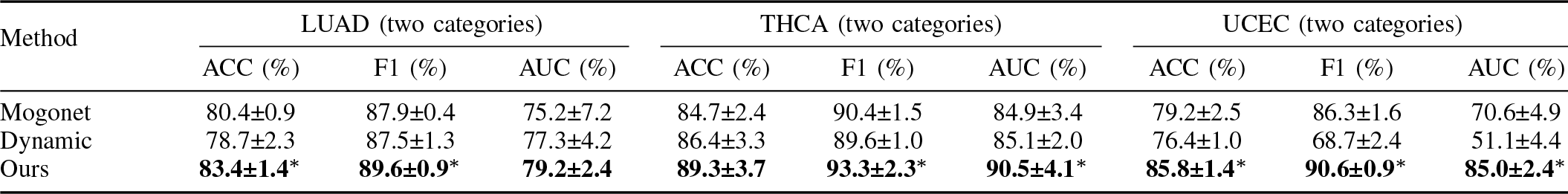
Comparison with state-of-the-art methods on the LUAD, THCA, and UCEC datasets. Mean and standard deviation are reported. The best results are in bold. Results with ‘*’ indicate that our model is significantly better (***P <*** 0.05) than the suboptimal method when using the independent t-test.

| Method | LUAD (two categories) |  |  | THCA (two categories) |  |  | UCEC (two categories) |  |  |
| --- | --- | --- | --- | --- | --- | --- | --- | --- | --- |
|  | ACC (%) | F1 (%) | AUC (%) | ACC (%) | F1 (%) | AUC (%) | ACC (%) | F1 (%) | AUC (%) |
| Mogonet | 80.4±0.9 | 87.9±0.4 | 75.2±7.2 | 84.7±2.4 | 90.4±1.5 | 84.9±3.4 | 79.2±2.5 | 86.3±1.6 | 70.6±4.9 |
| Dynamic | 78.7±2.3 | 87.5±1.3 | 77.3±4.2 | 86.4±3.3 | 89.6±1.0 | 85.1±2.0 | 76.4±1.0 | 68.7±2.4 | 51.1±4.4 |
| Ours | **83.4±1.4\*** | **89.6±0.9\*** | **79.2±2.4** | **89.3±3.7** | **93.3±2.3\*** | **90.5±4.1\*** | **85.8±1.4\*** | **90.6±0.9\*** | **85.0±2.4\*** |

### B. Ablation study

We conducted an ablation study to evaluate the efficacy of multi-level GAT (M-GAT) and TCP on four benchmark datasets. Specifically, we replaced the key components (M-GAT and/or TCP) with NN for comparison. As shown in Table V, both the M-GAT and TCP outperformed the simple neural networks. This demonstrates the effectiveness of these two components. M-GAT exhibits significant improvements over TCP on the ROSMAP and BRCA datasets, achieving an average improvement of 2.1% across three evaluation metrics, and also shows comparable performance with TCP for the LGG and KIPAN datasets. Our proposed model demonstrates the most promising performance across four datasets, underscoring the synergy effect of these two components.

**TABLE V.**
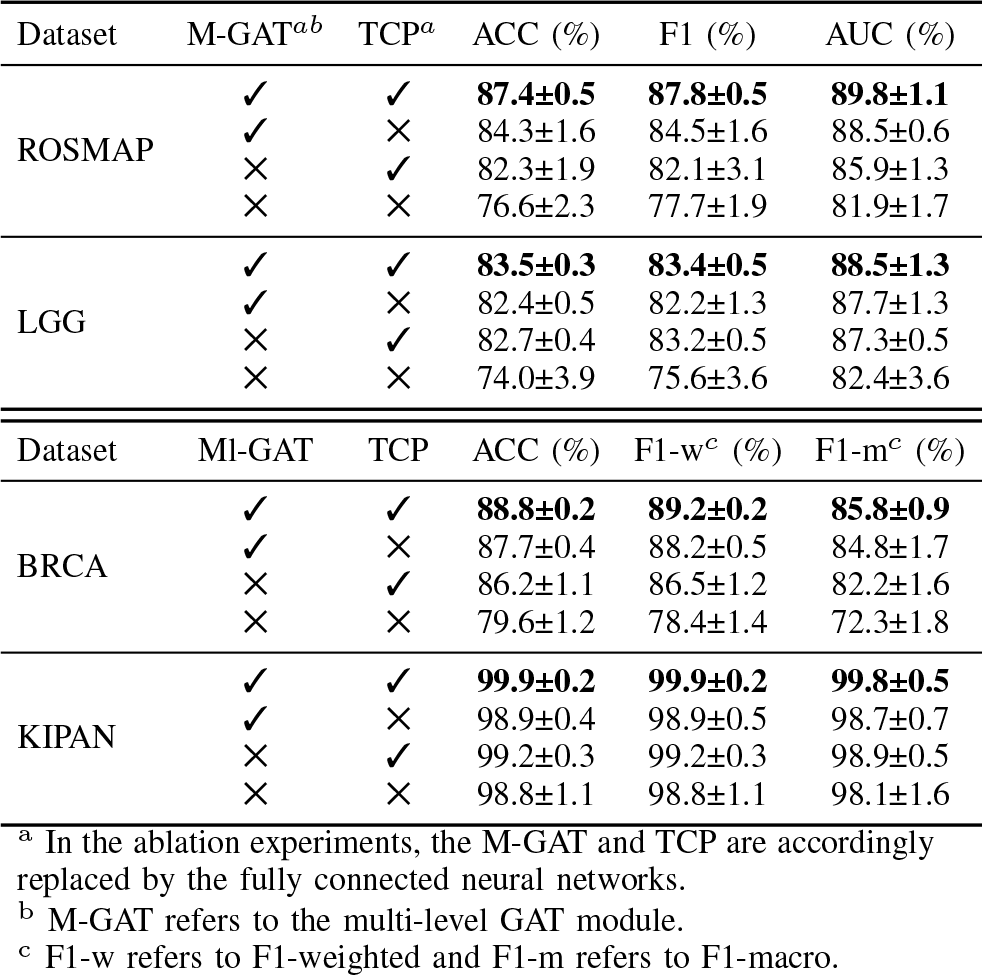
Ablation study of the key components in GREMI on benchmark datasets. The best results are in bold.

### C. Performance comparison under different omics types

To demonstrate that multi-omics could provide complementary information from multiple perspectives, we compared the classification performance when utilizing different types of omics data. Fig. 3 illustrated that the combination of all three omics outperformed other conditions (i.e., two and single omics data). This highlighted the unique contributions of each omics data, stressing the advantage of omics integration. Consequently, it can be deduced that GREMI boosts classification outcomes by fusing multi-omics, with improvements evident as more omics types are incorporated.

**Fig. 3.**
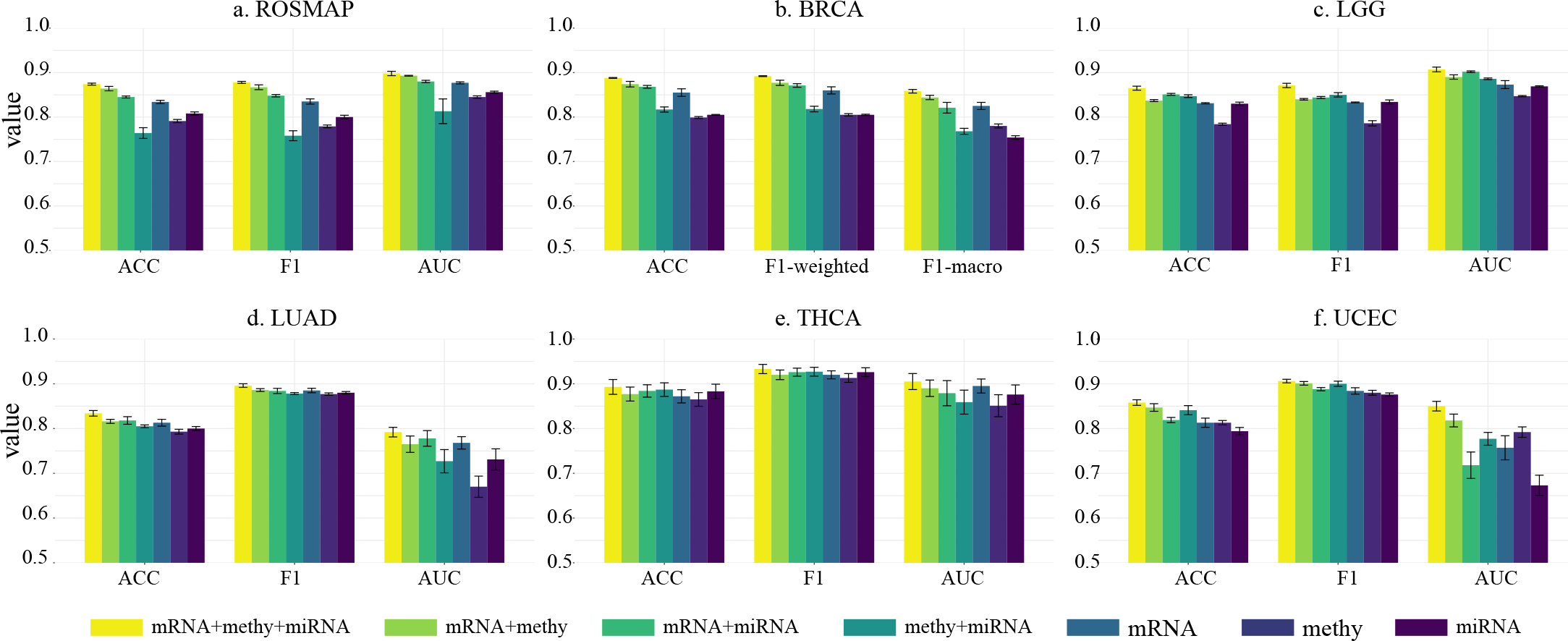
Performance comparison of the proposed method on different combinations of omics data. Means with standard errors are illustrated. Due to the easy classification of KIPAN, it is not presented here.

In certain tasks, our model using two modalities, outperformed Mogonet which utilized all three. For example, our mRNA+methy model surpassed Mogonet with three-omics in both ACC and F1 on ROSMAP (ACC: 86.4% *>* 81.5%, and F1: 86.7% *>* 82.1%). This was also observed in the BRCA subtype classification, where the GREMI mRNA+methy (ACC: 87.4%, F1-weighted: 87.7%) and mRNA+miRNA (ACC: 86.8%, F1-weighted: 87.1%) outperformed Mogonet using three modalities (ACC: 82.9%, F1-weighted: 82.5%). This further implied the effectiveness of sample-wise networks in characterizing features and the efficiency of our framework in information extraction and data fusion.

### D. Important modules identified by GREMI

For each omics, we generated a set of important modules specific to each sample and extracted consensus modules that existed across all samples. The details regarding the number of sample-wise modules and consensus modules are listed in Supplementary Table S1. Given the potential involvement of a limited number of molecules in biological regulation and considerations of computational complexity, we prioritized modules with sizes (i.e., number of nodes) ranging from two to five. For each disease, we assessed the statistical significance and functional relevance of top-ranked modules (i.e., modules with the highest importance scores). In addition, we evaluated the importance of individual biomarkers to provide a complementary explanation. From Table VI and Supplementary Results, we could observe that there were few overlaps between top individual biomarkers and top modules. This implied that the molecules did not always individually perform functions, but could potentially interact with others to jointly have an effect on diseases. Due to space limitations, we detailed only the modules identified from ROSMAP and BRCA in the subsequent sections, as shown in Table VI and Fig. IV. A detailed discussion of individual-level biomarkers of AD and BRCA and the top findings related to other diseases were available in the Supplementary Results.

**TABLE VI.**
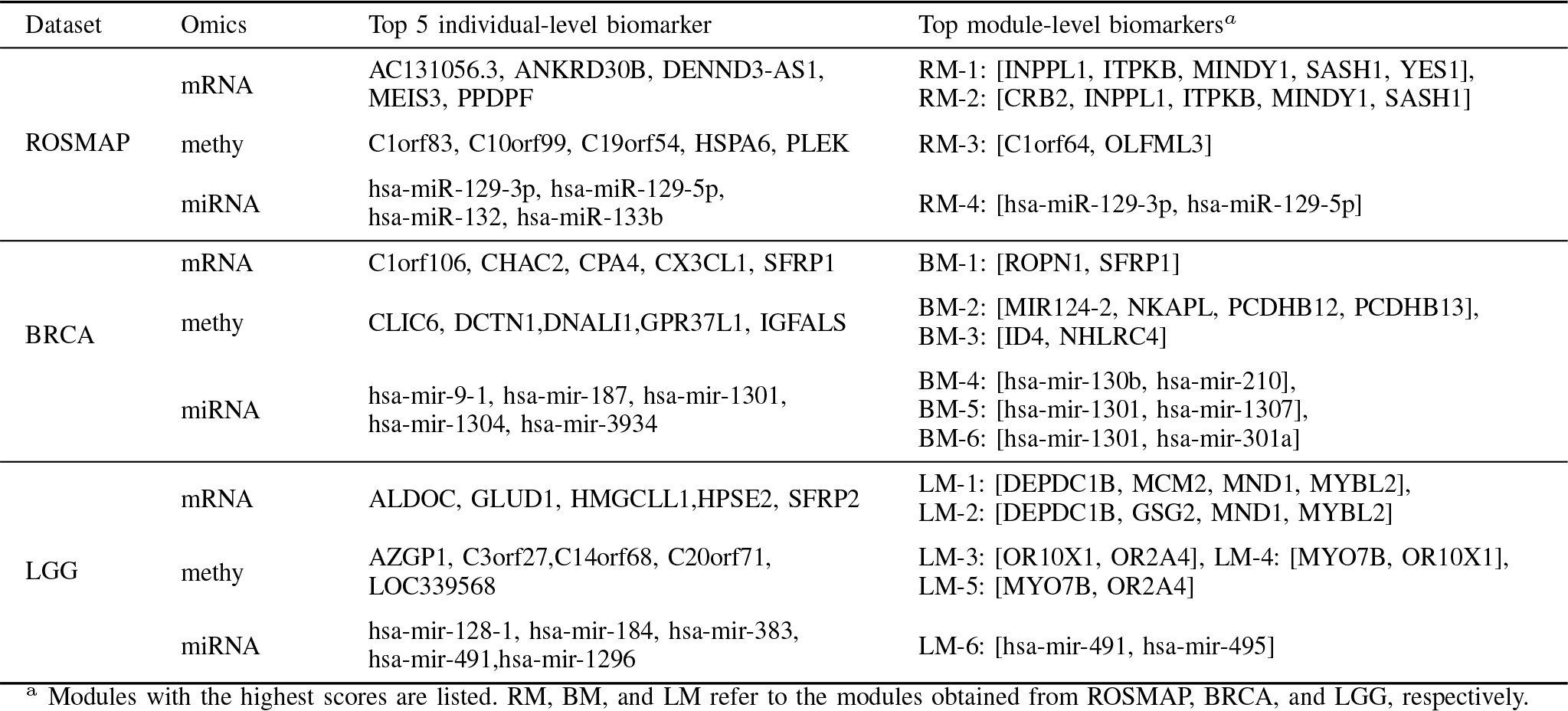
Top identified module-level and individual-level biomarkers.

**Fig. 4.**
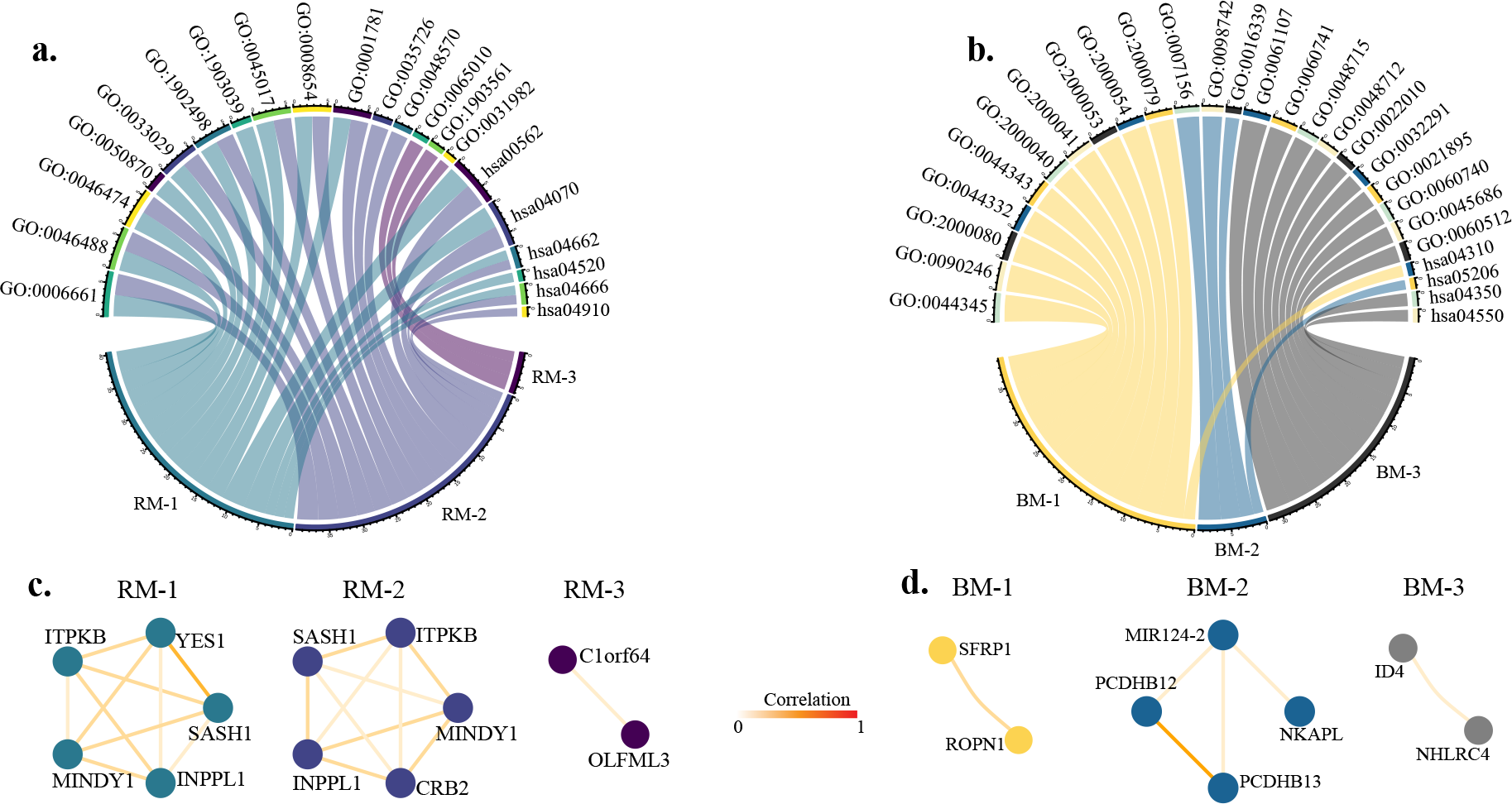
Functional annotation. a. Pathway enrichment of ROSMAP modules. b. Pathway enrichment of BRCA modules. c. Co-functionality of ROSMAP modules. d. Co-functionality of BRCA modules.

#### 1) GREMI identified biomarkers related to AD

Using the MCTS algorithm on mRNA, methylation, and miRNA co-functional networks, we obtained eighteen, one, and two AD-related consensus modules, respectively. Table VI presented the most important modules, including two mRNA co-expression (RM-1 and RM-2), one co-methylation (RM-3), and one miRNA co-expression (RM-4) modules. We also listed the top five individual biomarkers for each omics data type (see Table VI). Then we explored the functional relevance of identified modules and individual biomarkers with AD and its progression through permutation tests, functional enrichment, and an extensive literature review.

The permutation test indicated that the importance scores of the top modules were significantly higher than random subgraphs (i.e., all modules obtained permutation *P <* 1E-3). Fig. 4(a) shows the top enrichment results of AD modules, including the top ten GO terms and top five KEGG pathways (if available) enriched by each module. Additionally, we extracted and visualized the functional correlations among module elements (Fig. 4(c)). Given that one miRNA (hsa-miR-129-5p) was not available in both GO and KEGG databases, we did not perform functional annotation for RM-4.

There were several pathways enriched by both RM-1 and RM-2, given that four genes were commonly identified by these two modules. This suggested the crucial role and collective effect of these four molecules in driving biological processes relevant to disease pathogenesis. From Fig. 4, several GO terms related to the phosphatidylinositol process were enriched in RM-1 and RM-2, including phosphatidylinositol biosynthetic process (GO:0006661, *P* = 3.32E-2) and phosphatidylinositol metabolic process (GO:0046488, *P* = 3.32E-2). Emerging evidence suggested that A*β* oligomers impaired synaptic function in AD by interfering with the phosphoinositide signaling pathway, especially by reducing signaling through the Phosphatidylinositol Biphosphate signaling pathway. This highlighted a potential therapeutic strategy against A*β*-induced dysfunction [49], [50]. Identified KEGG pathways included inositol phosphate metabolism (hsa00562, *P* = 1.38E-3) and phosphatidylinositol signaling system (hsa04070, *P* = 1.38E-3). Protein tau hyperphosphorylation characterized AD. Alterations in inositol phosphate signaling influenced tau phosphorylation [51]. Among the enriched pathways of RM-3, extracellular vesicles (EVs, GO:1903561, *P* = 8.98E-3) presented as potential biomarkers for various diseases, including cancer and neurodegenerative diseases [52], [53]. EVs were implicated in tauopathy progression [54]. The presence of EVs in both plasma and cerebrospinal fluid further supported their potential as diagnostic markers [55].

#### 2) GREMI identified BRCA biomarkers

Table VI illustrates the top modules and biomarkers for BRCA. GREMI identified one mRNA, two methylation, and three miRNA co-functional modules. Fig. 4(b, d) shows the enriched pathways and structures of the top modules. Scores of the top modules were significantly higher than the randomly selected ones with permutation *P <* 1E-3. Wnt signaling-associated pathways, represented by GO:2000080 (*P* = 8.36E-3), GO:0044332 (*P* = 8.36E-3), and GO:0044343 (*P* = 8.36E-3), were predominant among the enriched pathways of BM-1. Wnt signaling pathway plays important roles in development and cancer progression, notably affecting breast cancer proliferation and metastasis [56]. Recent studies highlighted its influence on the immune microenvironment, stemness maintenance, therapeutic resistance, and breast cancer phenotypes [57]–[59]. In BM-2, pathways related to homophilic (GO:0007156, *P* = 2.82E-3) and calcium-dependent cell adhesion (GO:0098742, *P* = 3.87E-3) were significantly enriched, driven by plasma membrane adhesion molecules linked to breast cancer. Cell adhesion molecules (CAMs) were vital membrane receptors mediating interactions and intracellular signals essential for breast cancer processes. Exploring CAM signaling in breast cancer metastasis could reveal potential therapeutic targets [60]. Pathway enrichment of BM-3 suggested its link to breast cancer via the negative regulation of astrocyte differentiation. Astrocyte activation in triple-negative breast cancers (TNBCs) exacerbated malignancy and resistance to doxorubicin-induced apoptosis [61]. The aforementioned findings either directly affirmed or suggested associations between the identified modules and breast cancer.

### E. Validation of LGG biomarkers in an independent cohort

There were five genes identified from the top LGG mRNA modules, including *DEPDC1B, GSG2, MCM2, MND1*, and *MYBL2*. We validated these findings for their prediction ability using an independent mRNAseq dataset from the CGGA.

Given the potential effect of data imbalance, we employed the F1 score as the evaluation metric. With an SVM classifier, these five genes obtained an F1 score of 0.731 for LGG grade 2 versus grade 3 classification. To further evaluate the significance of classification performance, a permutation test was carried out 1,000 times, yielding a *P* of 4.8E-2. These confirmed both the statistical and practical significance of the identified LGG mRNA modules, emphasizing the robustness of our classification model and interpretation method.

## IV. Discussion and conclusions

The advancement of omics technologies and personalized medicine has yielded numerous supervised datasets valuable for predictive tasks, including disease diagnosis, tumor grading, and cancer subtyping. Effective integration of multi-omics data has proven its superiority over single omics in disease prediction. However, in order to make significant impacts in clinical practice, the integration models must offer accurate diagnostic guidance as well as encompass a broad spectrum of diseases, thereby highlighting the need for both high accuracy and robust generalization capability.

To achieve optimal multi-omics integration, several expectations should be considered. First, fully exploiting the information contained within each type of omics is essential to capture a comprehensive molecular characterization. Second, addressing the challenges posed by omics heterogeneity can significantly improve the performance of integration. Third, the inclusion of a broader range of omics types is anticipated to enhance the predictive capabilities.

In this context, we introduce GREMI, an innovative framework tailored to accommodate diverse omics data efficiently. The superior performance of GREMI is attributed to its comprehensive understanding of intra-omics interaction and adaptive utilization of inter-omics informativeness. Meanwhile, GREMI considered the multifactor-driven complexities of diseases. By combining omics data with topological structures, it established disease-specific networks incorporating functional interactions, reflecting recognition of contextualized molecular interactions in understanding disease mechanisms. Through true-class-probability-based fusion, GREMI adaptively utilized inter-omics informativeness, balancing prediction contributions of each omics to enhance both the precision and stability of the model. In our experiments, GREMI achieved state-of-the-art performance in all seven classification tasks. Furthermore, the ablation study confirmed the expectation that increasing the number of omics types could improve the molecular understanding of the disease. Through its implementation across seven different diseases, our model offers a more precise comprehension of disease mechanisms and patient categorization. The excellent performance and generalizability of GREMI highlight its potential for efficiently managing multiple complex diseases in practical clinical settings.

In addition to the accurate classification capabilities, GREMI identified important modules through a GNN interpretation from a co-functionality perspective. By uncovering disease-associated modules beyond individual genes, our explanation strategy aligned more closely with the multifactorial nature of complex diseases. The proposed local-global mixed strategy further promoted the extraction of subgraphs, highlighting the disease-relevant synergistic functions between molecules. The functional annotation and permutation test further demonstrated both the co-functionality and the disease-relevance of the identified module. Furthermore, an independent dataset confirmed the robustness and efficacy of our framework in pinpointing potential therapeutic targets.

However, some limitations should be acknowledged. First, although the co-functional networks offered valuable disease-specific insights, they might simultaneously miss the general molecular representations contained in conventional PPI networks. Secondly, subgraph exploration was relatively complex as the MCTS needed to evaluate each pruning action to determine the candidate ones. In our experiments, we capped the module size at five because it was time-consuming to generate a large number of random subgraphs with the same size in each permutation test. Given that the permutation evaluation was not required in practice, this parameter can be adjusted to produce larger candidate modules.

In the future, we plan to develop a hybrid knowledge-guided model that employs both disease-specific with generalized PPI networks, to deepen our understanding of multifactor-driven pathogenesis. Furthermore, stability and generalizability studies can be performed to investigate if the model trained from a specific disease can be transferred to various conditions.

## V. Supplementary materials

Supplementary information accompanies this paper at https://github.com/Yaolab-fantastic/GREMI.

## VI. Data availability

The raw ROSMAP omics data can be accessed at https://doi.org/10.7303/syn2580853. The raw omics data of BRCA, LGG, KIPAN, LUAD, THCA, and UCEC from The Cancer Genome Atlas are available at https://portal.gdc.cancer.gov/. The raw omics data of LGG-V from the Chinese Glioma Genome Atlas are accessible at http://www.cgga.org.cn/. All eight datasets used in our model training are available at https://github.com/Yaolab-fantastic/GREMI.

## Notes

This work is partly supported by the National Natural Science Foundation of China (62102115, 62103116, 82151303), the Shandong Provincial Natural Science Foundation (2022HWYQ-093), the Natural Science Foundation of Heilongjiang Province (LH2022F016), the Fundamental Research Funds for the Central Universities (3072022TS2614), the STI-2030-Major Projects (2021ZD0204002), and Key-Area Research and Development Program of Guangdong Province (2019B030335001).

### Competing Interest Statement

The authors have declared no competing interest.

### Summary of Updates

Updating the prediction phase and experiments. Adding a novel interpretation method and the results. Adding a validition part of the identified biomarkers.

